# Replicated chromatin curtails 53BP1 recruitment in BRCA1-proficient and -deficient cells

**DOI:** 10.1101/2020.02.24.947168

**Authors:** Jone Michelena, Stefania Pellegrino, Vincent Spegg, Matthias Altmeyer

## Abstract

DNA double-strand breaks can be repaired by two competing mechanisms, non-homologous end-joining (NHEJ) and homologous recombination (HR). Whether one or the other repair pathway is favored depends on the availability of an undamaged template DNA that allows for homology-directed repair. The tumor suppressor proteins 53BP1 and BRCA1 are considered antagonistic players in this repair pathway choice, as 53BP1 restrains DNA end resection, whereas BRCA1, together with its partner protein BARD1, displaces 53BP1 from damaged replicated chromatin and promotes HR. How cells switch from a 53BP1-dominated to a BRCA1-dominated response as they progress through the cell cycle is incompletely understood. Here we reveal, using high-throughput microscopy and applying single cell normalization to control for increased genome size as cells replicate their DNA, that 53BP1 recruitment to damaged replicated chromatin is inefficient in both BRCA1-proficient and BRCA1-deficient cells, in comparison to 53BP1 accumulation at damaged unreplicated chromatin. These findings substantiate a dual switch model from a 53BP1-dominated response in unreplicated chromatin to a BRCA1-BARD1-dominated response in replicated chromatin, in which replication-coupled dilution of 53BP1’s binding mark H4K20me2 functionally cooperates with BRCA1-BARD1-mediated suppression of 53BP1 binding. More generally, we suggest that appropriate normalization of single cell data, e.g. to DNA content, provides additional layers of information, which can be critical for quantifying and interpreting cellular phenotypes.

## Introduction

The balance between non-homologous end-joining (NHEJ) and homologous recombination (HR) has important implications for maintenance of genome stability, for exploiting DNA damage response (DDR) defects as vulnerabilities in cancer therapy, and for harnessing the full potential of CRISPR/Cas9-mediated genome editing and gene therapy^1-3^. The choice between NHEJ and HR is closely linked to the cell cycle and to whether or not an undamaged homologous stretch of DNA is available as a template for HR^4-6^. While 53BP1-RIF1-Shieldin restrains DNA end resection of DNA double-strand breaks (DSBs) in the absence of a homologous template strand, after DNA replication BRCA1-BARD1 counteracts 53BP1 binding to damaged chromatin and promotes HR reactions^7-12^. The antagonism between 53BP1-RIF1-Shieldin and BRCA1-BARD1 has important implications for cancer therapy, in particular for targeting BRCA-deficient tumors with PARP inhibitors, and for improving CRISPR/Cas9-mediated gene editing.

HR is confined to the S and G2 phase of the cell cycle, yet how the DDR discriminates between replicated and unreplicated areas of the genome during S phase progression has only started to emerge recently. Such discrimination is important for HR reactions to be favored when DSBs occur in replicated DNA, and mutagenic end resection to be prevented when DSBs occur in areas of the genome that were not replicated yet. Accumulating evidence suggests that this discrimination is linked to the different chromatin makeup ahead of and behind replication forks^10-14^. Of particular interest for the antagonism between the 53BP1 and BRCA1 protein complexes is the H4K20 dimethylation mark (H4K20me2), which is abundant in unreplicated chromatin on parental histones and absent from newly incorporated histones. Hence H4K20me2 is diluted in nascent replicated chromatin and only restored in mature chromatin in late G2/M^10-14^. 53BP1 binds H4K20me2 via its tandem tudor domain (TTD), and both H4K20me2 and the TTD are required for 53BP1 recruitment to DSBs^15, 16^. Conversely, unmethylated H4K20 (H4K20me0) in replicated chromatin is recognized by the HR-promoting protein complexes TONSL-MMS22L and BRCA1-BARD1^12, 14^.

Among the evolving methods to probe cellular stress responses at the single cell level is quantitative image-based cytometry (QIBC), which uses automated high-content microscopy in a cytometry-like fashion to stage individual cells according to their position in the cell cycle^17-19^. Besides quantification of multiple cellular parameters in large cell populations, when such microscopy-based single cell measurements are accurate enough to discriminate cells according to their position in the cell cycle, they allow for cell cycle-related data normalization, e.g. to the size of a cell or its nucleus, or to the replication status of the genome. As multiple cellular functions are tightly linked to cell cycle progression and to the associated duplication of cellular contents, including duplication of the genome and related entities, we suggest, using the example of cell cycle regulated 53BP1 recruitment to damaged chromatin as paradigm, that normalization of cell cycle-related phenotypes to DNA content and thus cell cycle position, when implemented in quantitative cell image analysis pipelines, can reveal additional layers of information that facilitate biological interpretation of quantitative cell imaging data.

It was recently established that the ankyrin repeat domain (ARD) of BARD1 binds to H4K20me0 and brings the BRCA1-BARD1 complex to DNA breaks when a sister chromatid is available for DSB repair by HR, thus illuminating the mechanistic underpinnings of BRCA1-mediated displacement of 53BP1 from replicated chromatin^12^. An unresolved question is, however, whether the antagonism between BRCA1-BARD1 and 53BP1 alone is sufficient to explain the displacement of 53BP1 from damaged replicated chromatin, or whether the dilution of H4K20me2 behind replication forks directly impacts 53BP1 recruitment and thereby affects its functions even when BRCA1-BARD1 functions are impaired. Elucidating the relative contribution of BRCA1-BARD1-related versus unrelated factors for antagonizing 53BP1’s functions could provide critical information on DSB repair pathway choice with implications for utilizing HR and NHEJ in the context of genome editing and for targeting the DDR in cancer therapy.

## Results and discussion

By means of software-assisted image segmentation and feature extraction, QIBC allows measurements of cellular phenotypes in large, asynchronously growing cell populations. Segmentation masks defined by the DAPI signal can be used to detect individual cell nuclei (Figure 1a). By multi-color imaging, additional cell cycle markers, such as the DNA replication marker 5-Ethynyl-2’-deoxyuridine (EdU) and Cyclin A (other useful markers include Cyclin B and the mitotic marker H3pS10), can be acquired in conjunction with a DDR marker of interest (Figure 1b). Validation of correct cell cycle staging at the level of individual cells occurs in both directions, from single cell images to the position of the cell in image-based cell cycle profiles, and conversely from any position in the one- or two-dimensional cell cycle profiles to the individual cell image (Figure 1c). Additional segmentation masks can be applied to detect sub-nuclear structures such as ionizing radiation (IR) induced foci (Figure 1d). By use of appropriate markers, phenotypes at any particular position in the cell cycle can thus be interrogated, without the need for cell cycle perturbations such as synchronization and without the need to categorize (i.e. gate) cells into binned groups. Of note, in such experiments the total DAPI intensity per nucleus scales linearly with DNA content, doubling as cells go from G1 (2N) to G2 (4N), and can therefore be used as a direct measure of genome size (Figure 1c).

**Figure 1:**
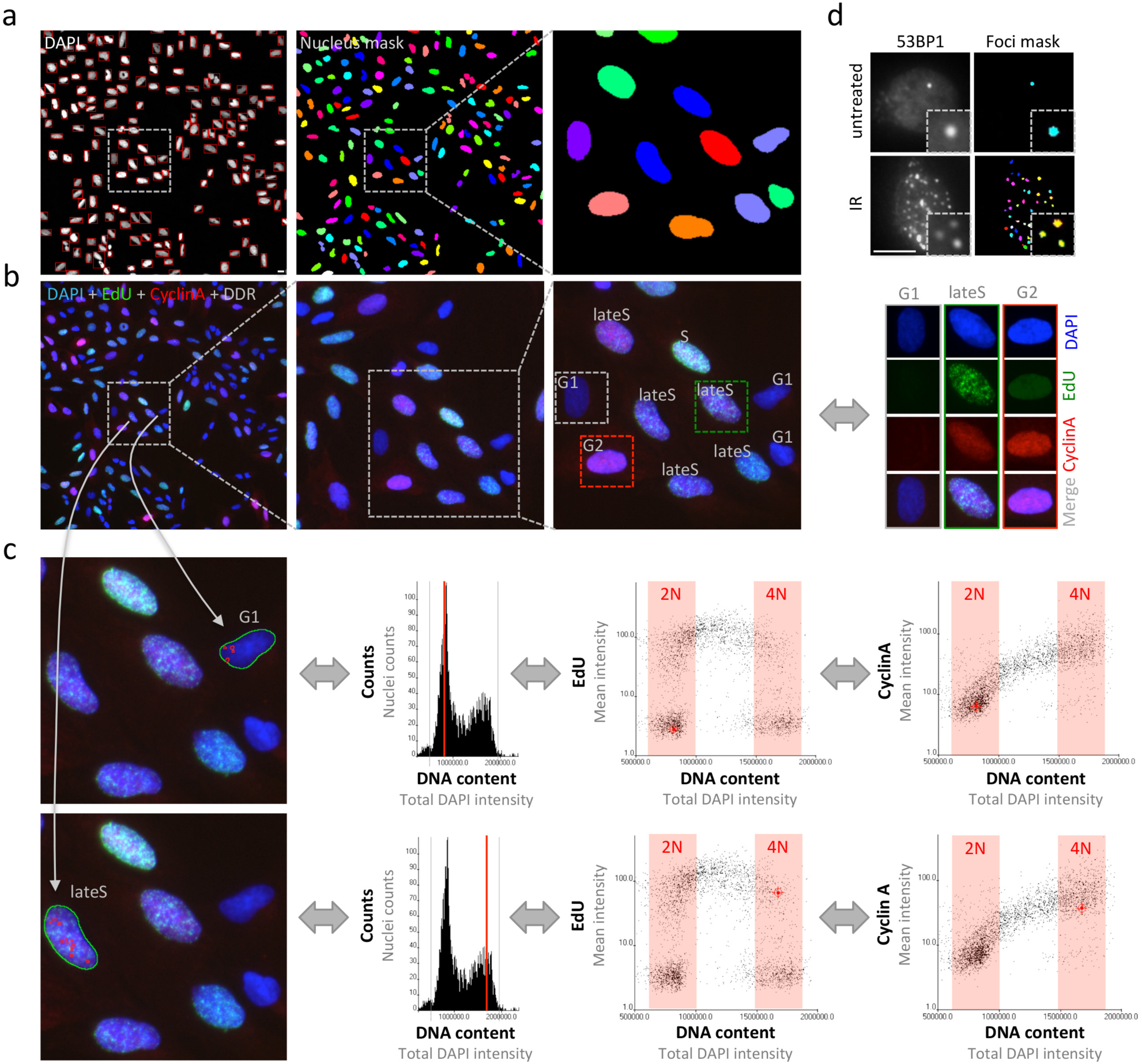
Cell cycle staging by QIBC. **(a)** Image segmentation based on the DAPI signal to detect individual cell nuclei. Typically, between 50 and 100 images per condition are acquired, yielding image information of between 3000 and 10000 cells. **(b)** In multi-color imaging experiments, appropriate cell cycle markers (here DAPI, EdU, Cyclin A) are combined with markers of interests to allow cell cycle-resolved interrogations of cellular responses. When multiple cell cycle markers are rationally combined and cross-compared, precise cell cycle staging can be achieved (e.g. to discriminate early G2 from late G2 cells or late G2 cell from mitotic cells). **(c)** Cell cycle profiles are generated for each cell population (here a one-dimensional cell cycle profile based on DAPI as well as two-dimensional cell cycle profiles based on EdU versus DAPI and Cyclin A versus DAPI). In the upper micrograph, a single cell in G1 is selected and its cell cycle position in the cell cycle profiles to the right is indicated in red. In the lower micrograph, a cell in late S-phase is selected and its cell cycle position in the cell cycle profiles to the right is indicated in red. As images and numerical data are linked, the analysis works in both ways, from single cell image to cell cycle profile, and reciprocally from cell cycle profile to single cell images, allowing for both quantification-based and image-based explorations of cell cycle related phenotypes. **(d)** For each individual cell nucleus, sub-nuclear structures such as IR-induced foci (here 53BP1) are segmented and their number and signal intensities are quantified to yield cell cycle-resolved maps of DNA damage responses. Scale bars, 10μm.

Scoring IR-induced 53BP1 foci by QIBC, we previously reported a gradual decline in 53BP1 accumulation at damaged chromatin as cells go through S-phase^11^, consistent with other studies^9, 10^. IR-induced DNA damage scales with the amount of DNA present in a cell’s nucleus and doubles as cells replicate their genome (Figure 2a). Consistently, when we measured DNA content and γH2AX foci formation (15 minutes after IR) as marker of IR-induced DSBs in a cell cycle resolved manner, both DNA content and γH2AX foci approximately doubled as cells went from G1 to G2 (Figure 2b, c). This is in agreement with prior work on irradiation-induced DNA damage load scaling linearly with the amount of DNA exposed to IR^20-24^, and with standard normalization procedures when working with cell populations synchronized at different phases of the cell cycle (e.g. for DSB measurements by pulse field gel electrophoresis (PFGE) twice as many G1-arrested cells as G2-arrested cells are loaded on the gel to correct for the difference in DNA content^24^). We thus decided to take a closer look at the recruitment of 53BP1 in presence or absence of its antagonist BRCA1-BARD1, taking into account that the induced DNA damage is proportional to the amount of DNA present in the nucleus. As expected, single cell normalization to DNA content in BRCA1-BARD1-proficient cells revealed a pronounced decline in 53BP1 foci formation as cells go through S-phase, both in antibody staining-based detection of endogenous 53BP1 (Extended Data Figures 1a-d and 2a-d), and in CRISPR/Cas9-engineered cells expressing fluorescently labeled 53BP1 from its natural gene promoter (Figure 3a-e). Importantly, single cell normalization to DNA content also revealed that IR-induced 53BP1 foci formation gradually declined as a function of DNA replication upon depletion of either BRCA1 or BARD1, although to a lesser extent as in BRCA1-BARD1-proficient cells (Figure 4a-c and Extended Data Figures 3a-d and 4a-b). The delta in efficiency of 53BP1 recruitment in gated cell subpopulation averages based on DNA content (2N versus 4N) was even more pronounced when G1 cells were compared to cells in late S / early G2 based on DAPI and Cyclin A staining, both in BRCA1-BARD1-proficient and -deficient cells (Figure 5a-d). Similarly, when H4K20me2 was used to define G1 versus late S / early G2 cells (i.e. excluding cells in mid and late G2 with gradually restored H4K20me2), the difference in 53BP1 recruitment was more pronounced compared to cells gated merely based on their DNA content (Figure 6a-d). Consistent results were obtained with the breast cancer cell line SUM149PT carrying the *BRCA1* 2288delT mutation and allelic *BRCA1* loss (Extended Data Figure 5a-c). Taken together, we conclude that inefficient 53BP1 recruitment to damaged replicated chromatin is an inherent feature that occurs both in BRCA1-deficient and, in a more pronounced manner, in BRCA1-proficient cells, and we therefore suggest that replication-coupled dilution of H4K20me2, in addition to enabling H4K20me0-mediated BRCA1-BARD1 recruitment, also directly affects the efficiency of 53BP1 recruitment in response to DNA damage.

**Figure 2:**
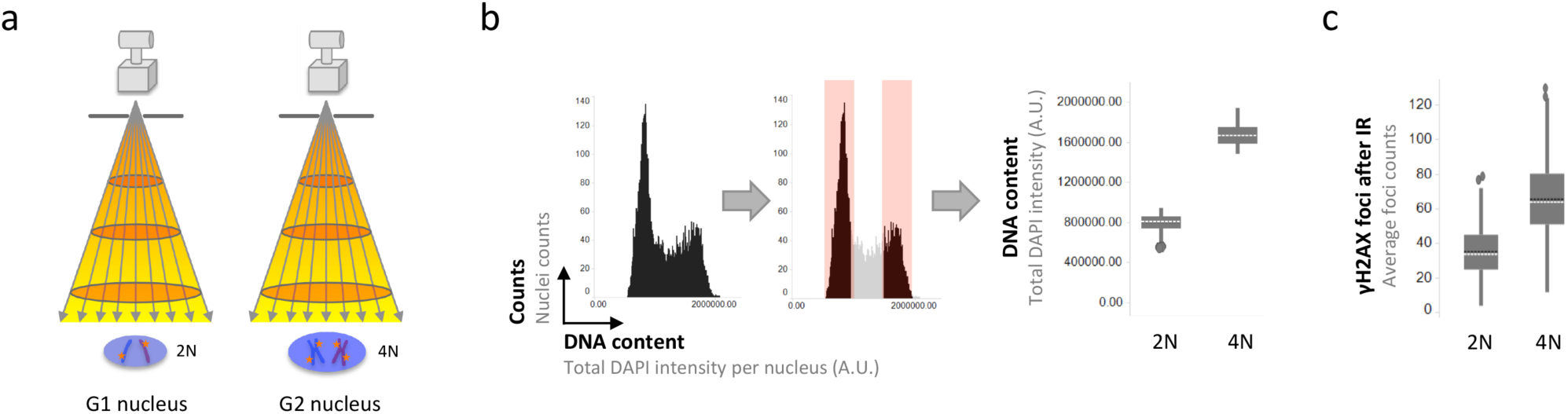
IR-induced DNA damage scales with DNA content. **(a)** Scheme to illustrate that ionizing radiation (IR) induced DNA damage scales with genome size. Due to the increased genome size when comparing G2 cells to G1 cells, more DNA damage occurs in an irradiated G2 cell nucleus compared to a G1 cell nucleus. **(b)** Quantitative image-based cytometry (QIBC) allows for cell cycle profiling based on the DAPI signal as proxy of DNA content. Accordingly, the DAPI signal in U-2 OS cells with a 4N DNA content (G2) is twice as high as the DAPI signal in cells with a 2N DNA content (G1). **(c)** Consistently, approximately twice as many γH2AX foci, as marker of DNA damage, are quantified in G2 cells as compared to G1 cells at early time-points after IR. U-2 OS cells were treated with 0.5 Gy of IR, fixed 15 minutes later, stained for DNA content and γH2AX, and γH2AX foci were quantified in a cell cycle-resolved manner by QIBC. Box plots with medians are shown.

**Figure 3:**
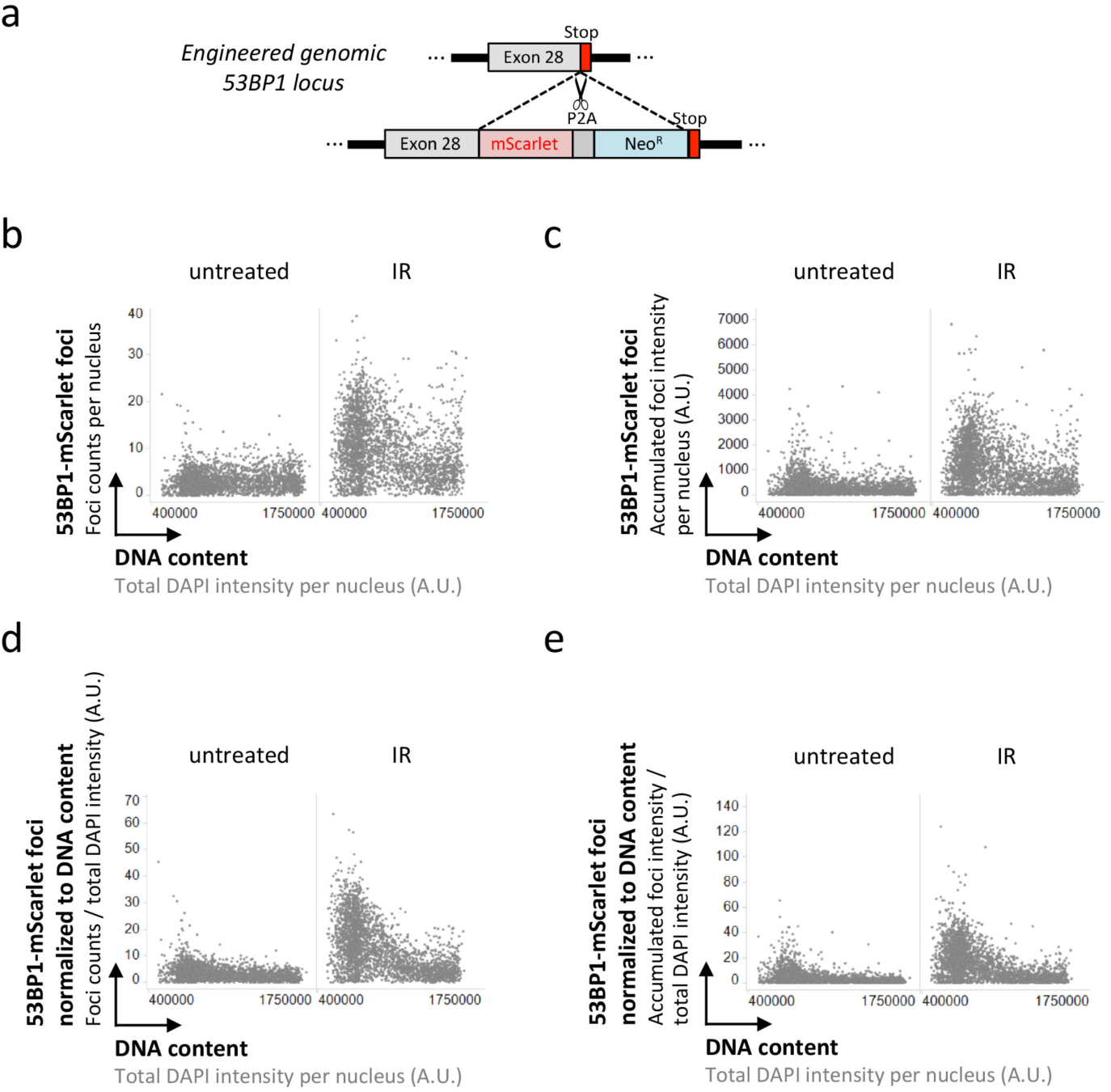
53BP1 response to IR-induced DNA damage as a function of cell cycle progression. **(a)** Scheme of the CRISPR/Cas9-based targeting of the endogenous 53BP1 gene locus to introduce mScarlet and generate cell lines expressing fluorescently labeled 53BP1 from its natural promoter. **(b)** U-2 OS 53BP1-mScarlet cells were treated with 0.5 Gy of IR, fixed 45 minutes later, stained for DNA content, and 53BP1-mScarlet foci counts were quantified by QIBC. **(c)** For the same cells shown in (b) the accumulated intensity of 53BP1-mScarlet foci per nucleus is shown. **(d)** The quantification from (b) was normalized at the single cell level to DNA content to control for increasing damage load with increasing DNA amount (see Figure 2). The parameter resulting from this normalization has arbitrary units and was multiplied by a multiple of 10 to yield data that could be plotted on linear scale in the depicted range. **(e)** For the same cells shown in (b), the accumulated intensity of 53BP1-mScarlet foci per nucleus was normalized to the DNA content of the same nucleus.

**Figure 4:**
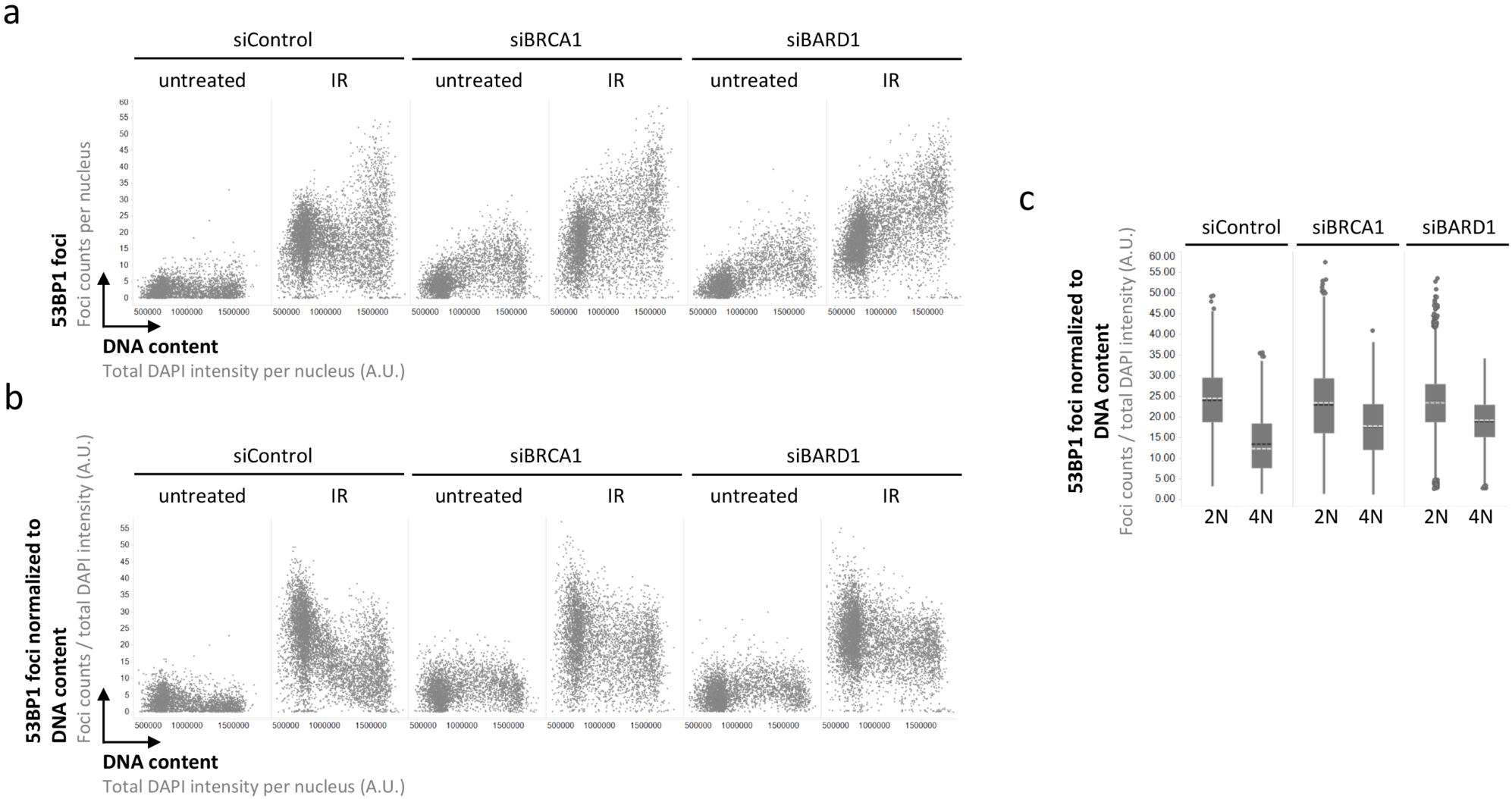
53BP1 recruitment to replicated chromatin is inefficient in absence of BRCA1-BARD1. **(a)** Cell cycle-resolved analysis of IR-induced 53BP1 foci formation in control conditions and upon depletion of either BRCA1 or BARD1. U-2 OS cells were treated with 0.5 Gy of IR, fixed 45 minutes later, stained for DNA content and 53BP1, and 53BP1 foci were quantified by QIBC. **(b)** The quantification from (a) was normalized at the single cell level to DNA content to control for increasing damage load with increasing DNA amount (see Figure 2). **(c)** Averaged relative 53BP1 recruitment after IR from (b) is compared in binned 2N versus 4N cells in presence or absence of BRCA1-BARD1. Box plots with medians are shown.

**Figure 5:**
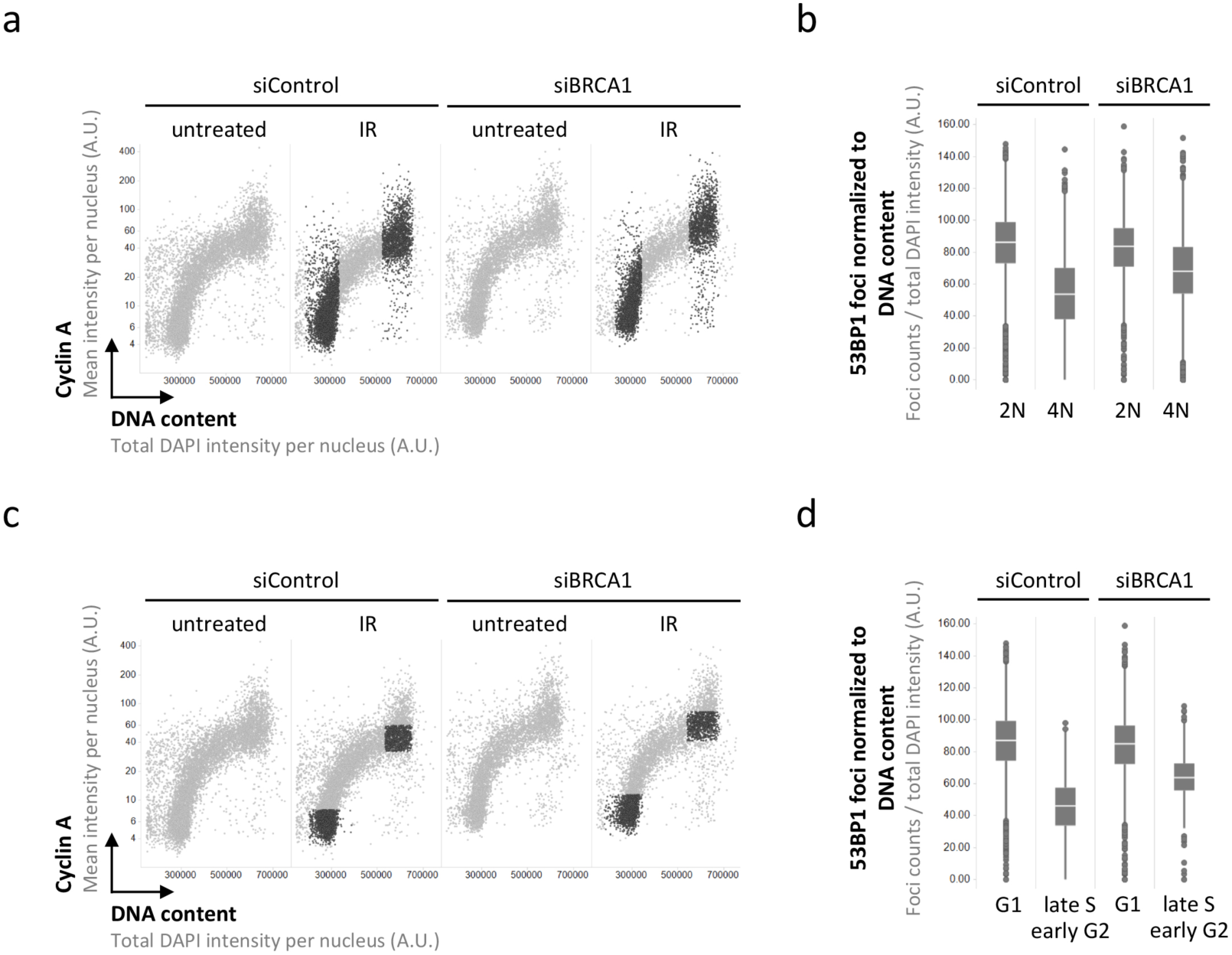
QIBC-assisted cell cycle gating based on DAPI and Cyclin A substantiates impaired 53BP1 recruitment to damaged replicated chromatin in both BRCA1-proficient and -deficient cells. **(a)** 2-D cell cycle profiles based on DAPI and Cyclin A. Cells with a 2N and 4N DNA content in the IR-treated samples marked in black. **(b)** Averaged relative 53BP1 recruitment after IR from the cells marked in black in (a) is compared. Box plots with medians are shown. **(c)** 2-D cell cycle profiles based on DAPI and Cyclin A. Cells in G1 and cells in late S / early G2 in the IR-treated samples marked in black. **(d)** Averaged relative 53BP1 recruitment after IR from the cells marked in black (c) is compared. Box plots with medians are shown.

**Figure 6:**
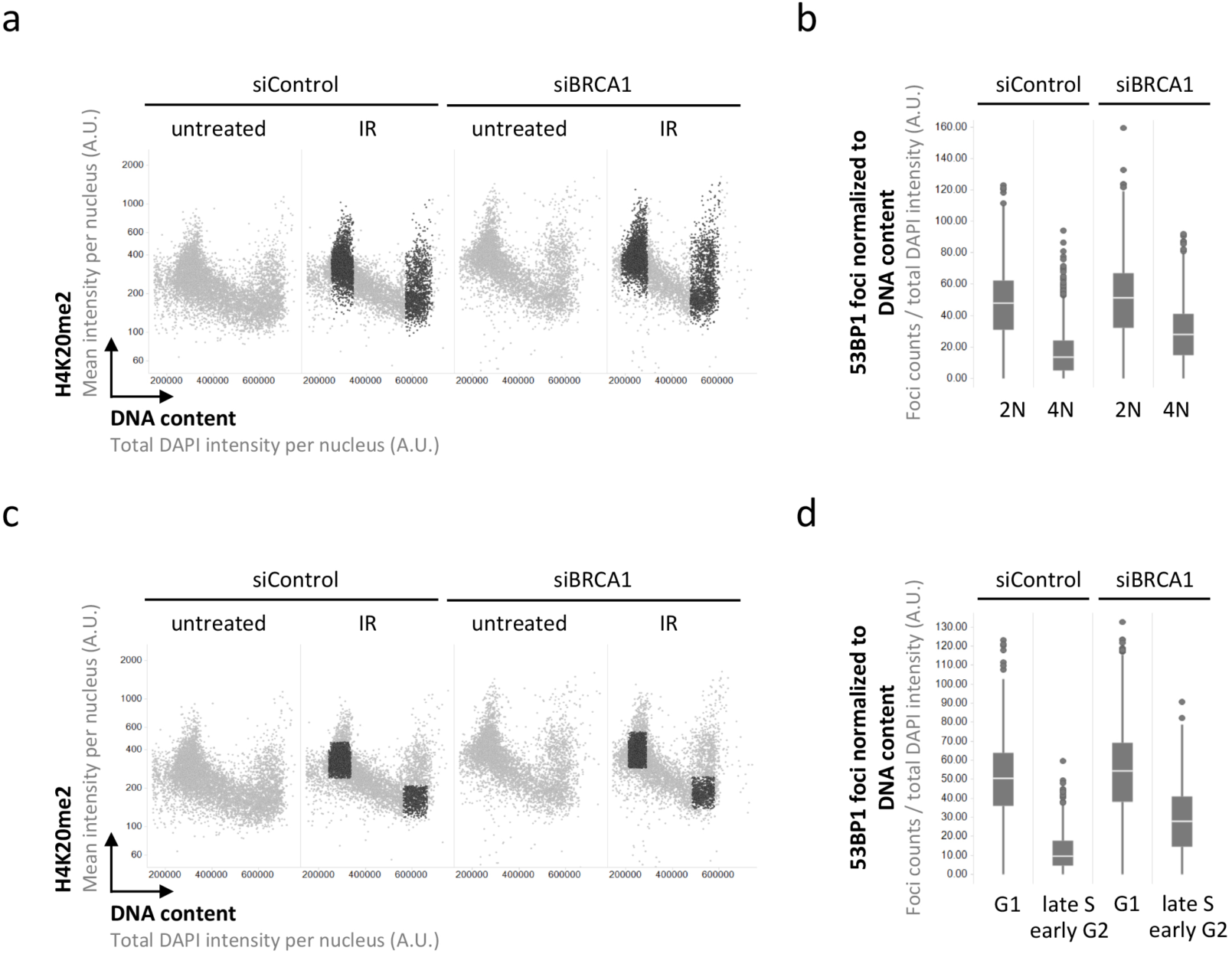
QIBC-assisted cell cycle gating based on DAPI and H4K20me2 substantiates impaired 53BP1 recruitment to damaged replicated chromatin in both BRCA1-proficient and -deficient cells. **(a)** 2-D cell cycle profiles based on DAPI and H4K20me2. Cells with a 2N and 4N DNA content in the IR-treated samples marked in black. **(b)** Averaged relative 53BP1 recruitment after IR from the cells marked in black in (a) is compared. Box plots with medians are shown. **(c)** 2-D cell cycle profiles based on DAPI and H4K20me2. Cells in G1 and cells in late S / early G2 in the IR-treated samples marked in black. **(d)** Averaged relative 53BP1 recruitment after IR from the cells marked in black (c) is compared. Box plots with medians are shown.

53BP1 requires its oligomerization domain for recruitment to sites of DNA damage and shows hallmarks of dynamic self-assembly by phase separation^25-27^. A reduced concentration of H4K20me2, as present in replicated nascent chromatin, therefore likely increases the threshold for efficient 53BP1 accumulation. Conversely, the high concentration of H4K20me2 in unreplicated chromatin (i.e. in G1/G0 cells and in unreplicated regions of the genome during S-phase progression), together with DNA damage-induced chromatin modifications that promote multivalent 53BP1 chromatin binding, provides a scaffold for efficient 53BP1 assembly around DNA break sites in the absence of a replicated template DNA required for HR repair. We therefore suggest that the effect of H4K20me2 dilution on 53BP1 assembly in nascent replicated chromatin functionally cooperates with the effect of H4K20me0-mediated BRCA1-BARD1 recruitment and the ensuing 53BP1 displacement, and that together they represent a dual switch to ensure that DSBs in unreplicated areas of the genome are protected from excessive DNA end resection and illegitimate recombination^28^ and channeled towards NHEJ, while DSBs in replicated areas of the genome are released from the DNA end protection functions of 53BP1 and channeled towards resection and HR (Figure 7). Accordingly, the mutual antagonism between 53BP1 and BRCA1-BARD1 may be seen as a bistable system, the robustness of which is achieved by cooperative effects resulting from high affinity 53BP1 and low affinity BRCA1-BARD1 binding to unreplicated chromatin versus low affinity 53BP1 and high affinity BRCA1-BARD1 binding in replicated chromatin. This view is consistent with supraphysiological 53BP1 accumulation at damaged replicated chromatin in BRCA1-deficient cells^29^, yet it suggests that even in absence of BRCA1-BARD1 the accumulation of 53BP1 is curtailed by reduced H4K20me2 in replicated chromatin. The DDR thus makes use of replication-coupled dilution of an abundant histone mark, which cells only restore in replicated chromatin when genome duplication has been completed, and enforced premature restoration of H4K20me2 during S-phase progression indeed shifts the balance towards a 53BP1-governed response^11^. In light of the growing interest to target the balance between HR and NHEJ in cancer therapy and to modify the underlying mechanisms for improved genome editing^30^, we envision the dual switch mechanism from an H4K20me2-53BP1-dominated response in unreplicated chromatin to an H4K20me0-BRCA1-BARD1-dominated response in replicated nascent chromatin to be of relevance.

**Figure 7:**
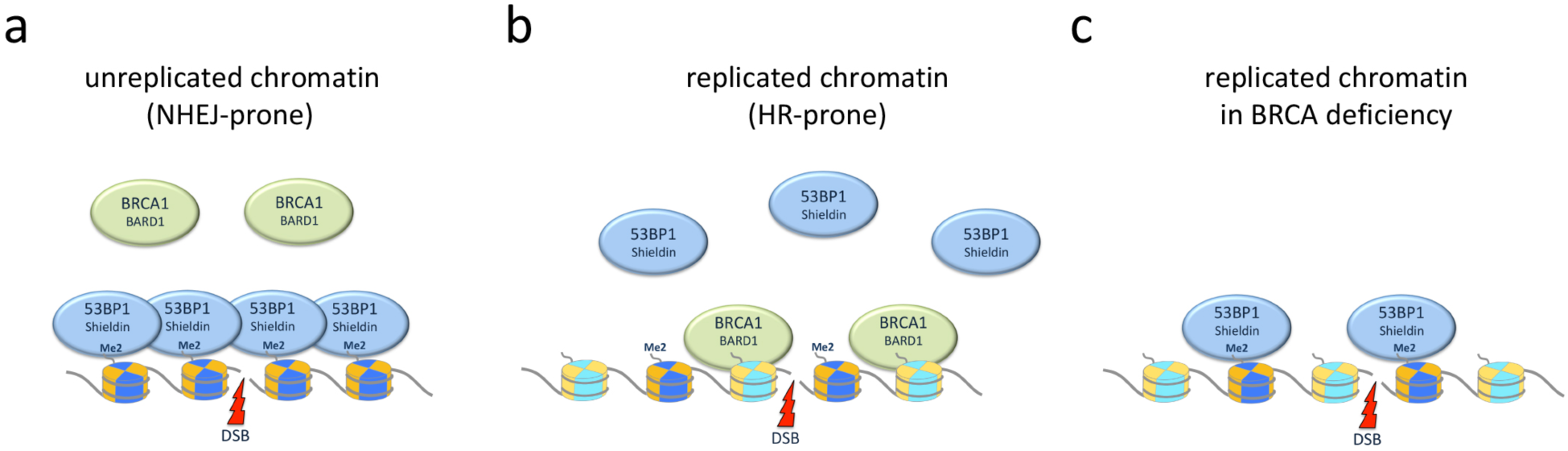
Simplified model of a dual switch to regulate the accumulation of 53BP1 and BRCA1-BARD1 at unreplicated versus replicated damaged chromatin. **(a)** The high density of H4K20me2 in unreplicated chromatin (in G1/G0, and throughout S-phase in yet to be replicated regions of the genome), together with DNA damage induced γH2AX and H2AK15ub (not shown), promotes efficient assembly of 53BP1 and its downstream effectors. This shields DSBs against excessive DNA end resection and illegitimate recombination and generally renders break sites NHEJ-prone. **(b)** As a dual switch, reduced 53BP1 binding upon replication-coupled dilution of H4K20me2 functionally cooperates with H4K20me0-mediated BRCA1-BARD1 recruitment to promote DNA end resection and HR reactions in replicated areas of the genome (during S-phase progression in nascent chromatin and prior to H4K20me2 restoration in late G2/M). **(c)** In BRCA1-BARD1-deficient cells, 53BP1 can accumulate at damaged replicated chromatin and exert some of its functions, however compared to unreplicated chromatin this recruitment is less efficient, likely reflecting the reduced density of H4K20me2.

More generally, we propose that, depending on the biological question, appropriate normalization becomes inevitable to interpret high-content single cell data, and that image-based normalization to cell size, nuclear volume, DNA content, or other suitable cell cycle markers can provide additional layers of information, which may be critical for quantifying and interpreting cellular responses to stress, including genotoxic stress by irradiation, chemotherapy, or newly emerging anti-cancer drugs.

## Extended Data Figures

**Extended Data Figure 1:**
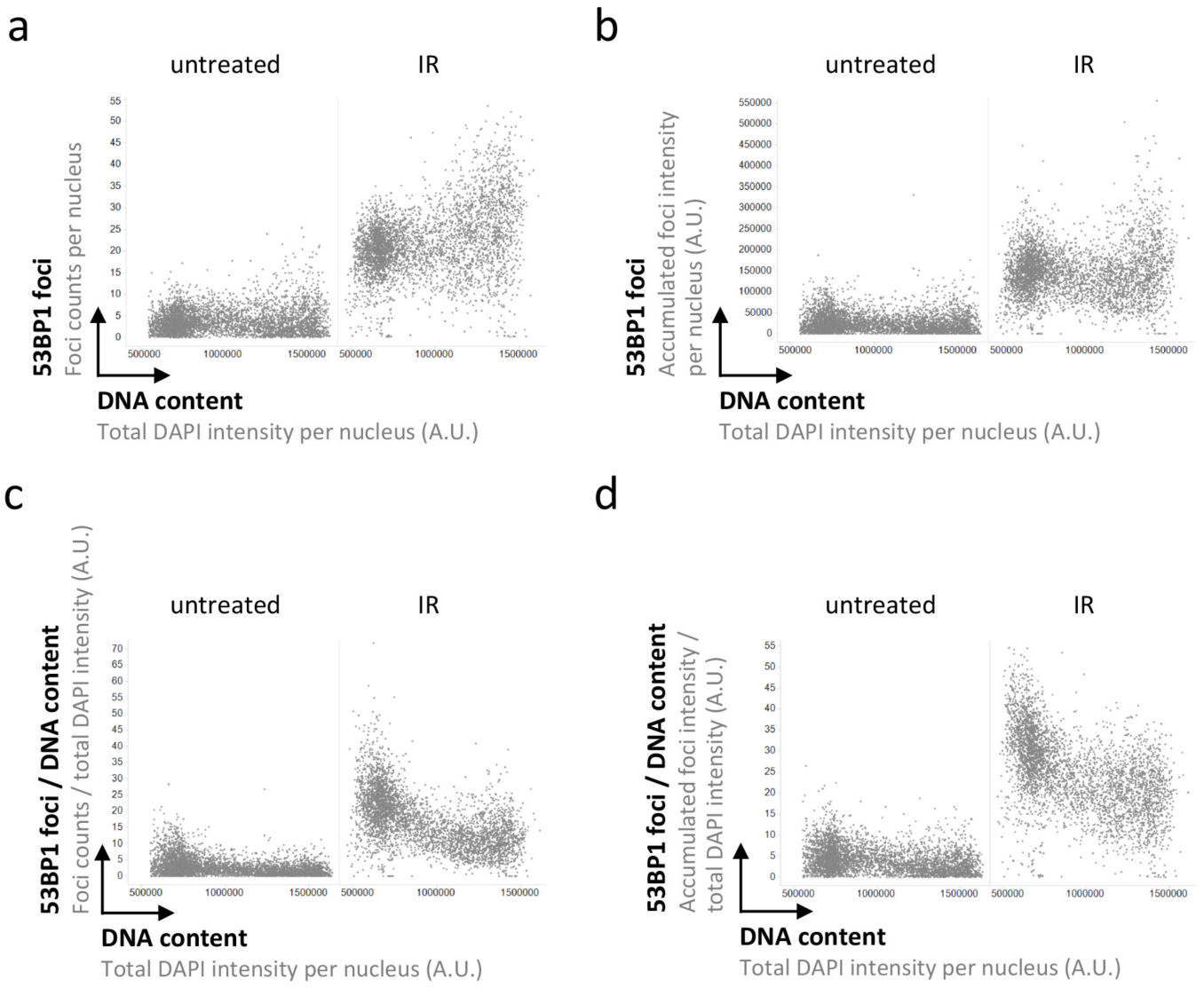
53BP1 response to IR-induced DNA damage (0.5 Gy) as a function of cell cycle progression. **(a)** U-2 OS cells were treated with 0.5 Gy of IR, fixed 45 minutes later, stained for DNA content and 53BP1, and 53BP1 foci counts were quantified by QIBC. **(b)** For the same cells shown in (a) the accumulated intensity of 53BP1 foci per nucleus is shown. **(c)** For the same cells shown in (a), the foci counts per nucleus were normalized to the DNA content of the same nucleus. **(d)** For the same cells shown in (a), the accumulated intensity of 53BP1 foci per nucleus was normalized to the DNA content of the same nucleus.

**Extended Data Figure 2:**
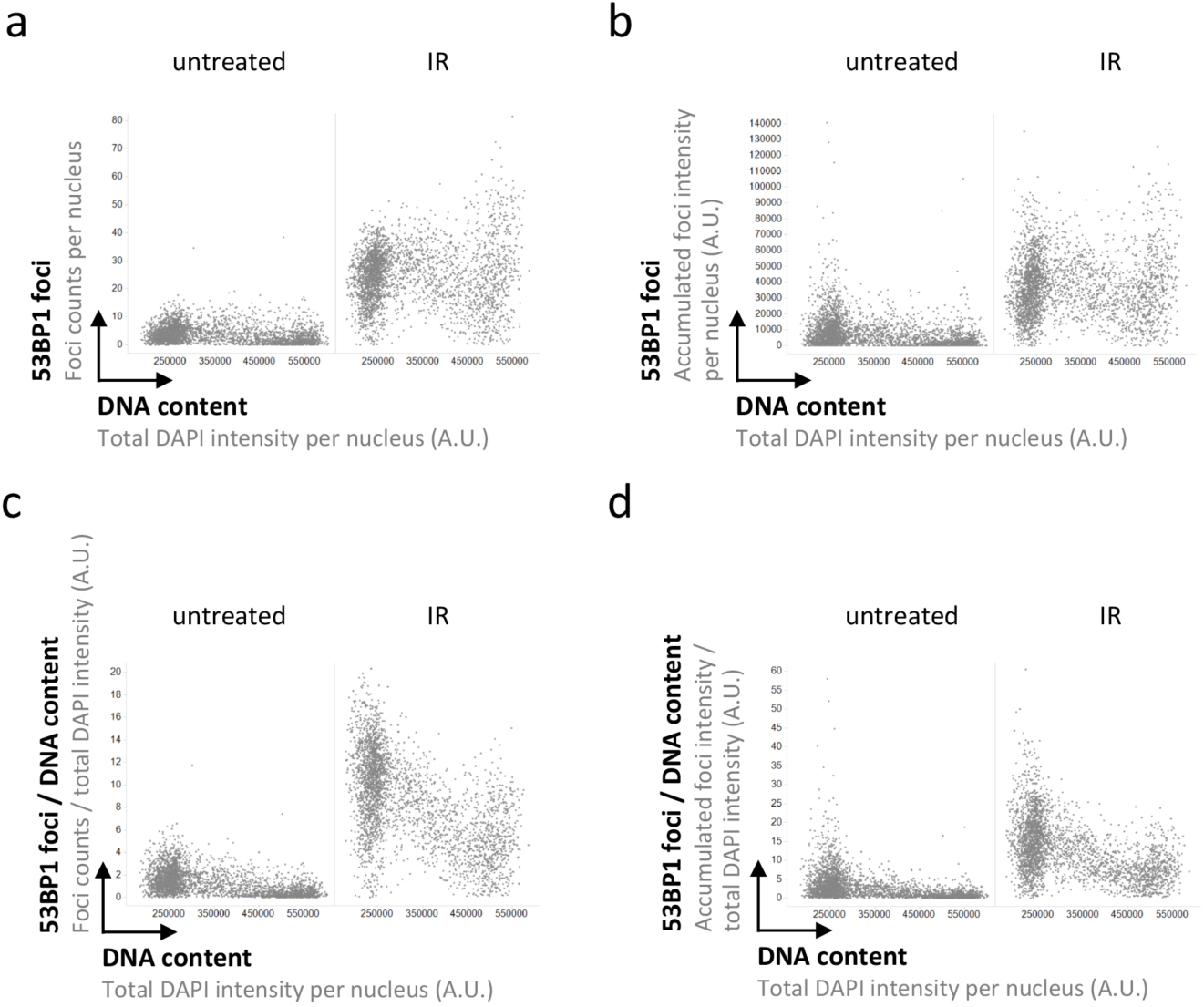
53BP1 response to IR-induced DNA damage (2 Gy) as a function of cell cycle progression. **(a)** U-2 OS cells were treated with 2 Gy of IR, fixed 2 hours later, stained for DNA content and 53BP1, and 53BP1 foci counts were quantified by QIBC. **(b)** For the same cells shown in (a) the accumulated intensity of 53BP1 foci per nucleus is shown. **(c)** For the same cells shown in (a), the foci counts per nucleus were normalized to the DNA content of the same nucleus. **(d)** For the same cells shown in (a), the accumulated intensity of 53BP1 foci per nucleus was normalized to the DNA content of the same nucleus.

**Extended Data Figure 3:**
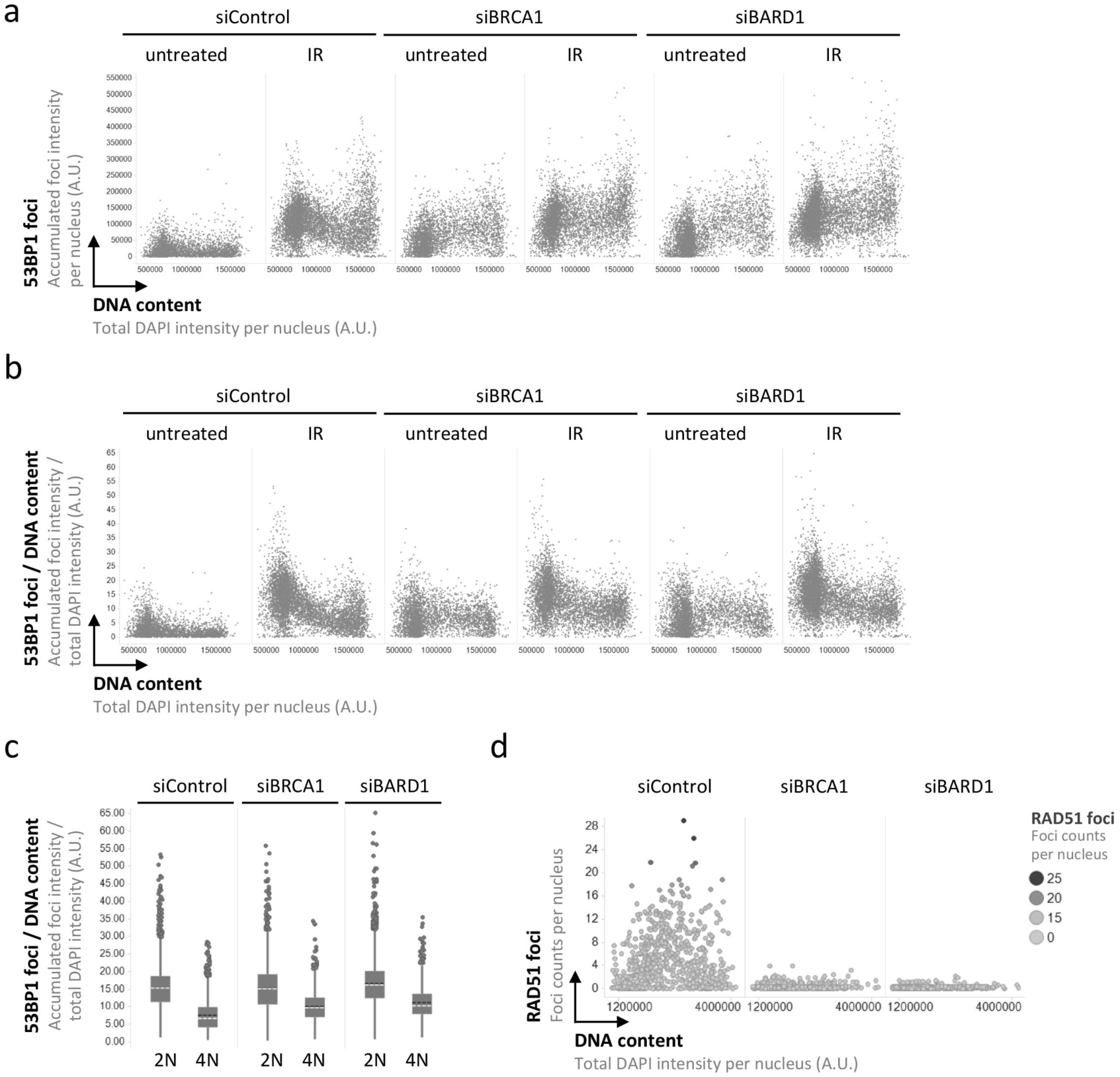
53BP1 recruitment to replicated chromatin is inefficient in absence of BRCA1-BARD1. **(a)** Accumulated intensity of 53BP1 foci per nucleus for the cells shown in Figure 4a. **(b)** Accumulated intensity of 53BP1 foci per nucleus normalized to the DNA content of the same nucleus for the cells shown in Figure 4. **(c)** Accumulated intensity of 53BP1 foci per nucleus normalized to the DNA content from (b) is compared in 2N versus 4N cells in presence or absence of BRCA1-BARD1. Box plots with medians are shown. **(d)** As control for the knockdown efficiency of BRCA1 and BARD1, nuclear RAD51 foci were analyzed by QIBC.

**Extended Data Figure 4:**
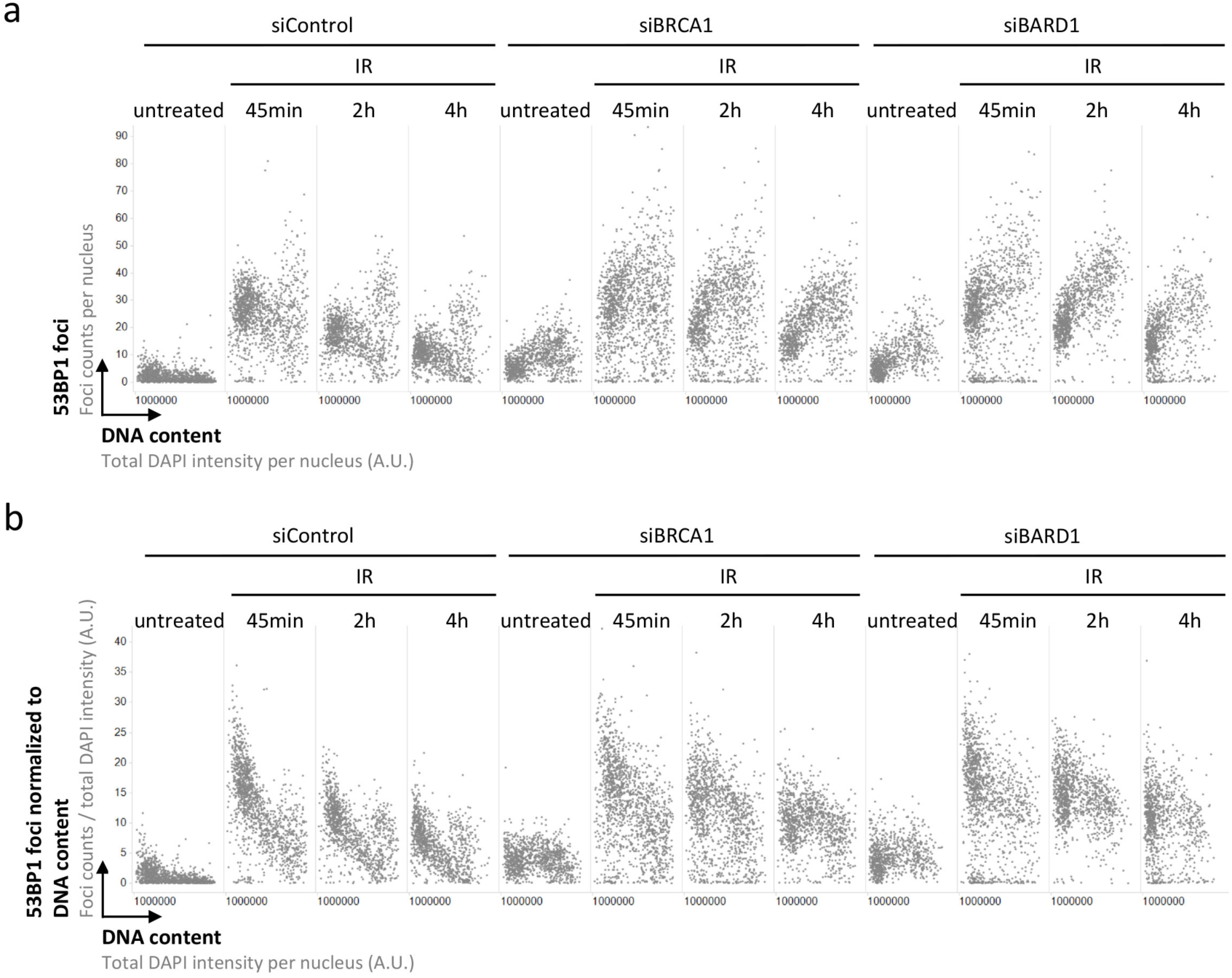
Extended dataset demonstrating that 53BP1 recruitment to replicated chromatin is inefficient in absence of BRCA1-BARD1. **(a)** Cell cycle-resolved analysis of IR-induced 53BP1 foci formation in control conditions and upon depletion of either BRCA1 or BARD1. U-2 OS cells were treated with 0.5 Gy of IR, fixed 45 minutes, 2h or 4h later, stained for DNA content and 53BP1, and 53BP1 foci were quantified by QIBC. **(b)** The quantification from (a) was normalized at the single cell level to DNA content to control for increasing damage load with increasing DNA amount. Corresponds to the data shown in Figure 4.

**Extended Data Figure 5:**
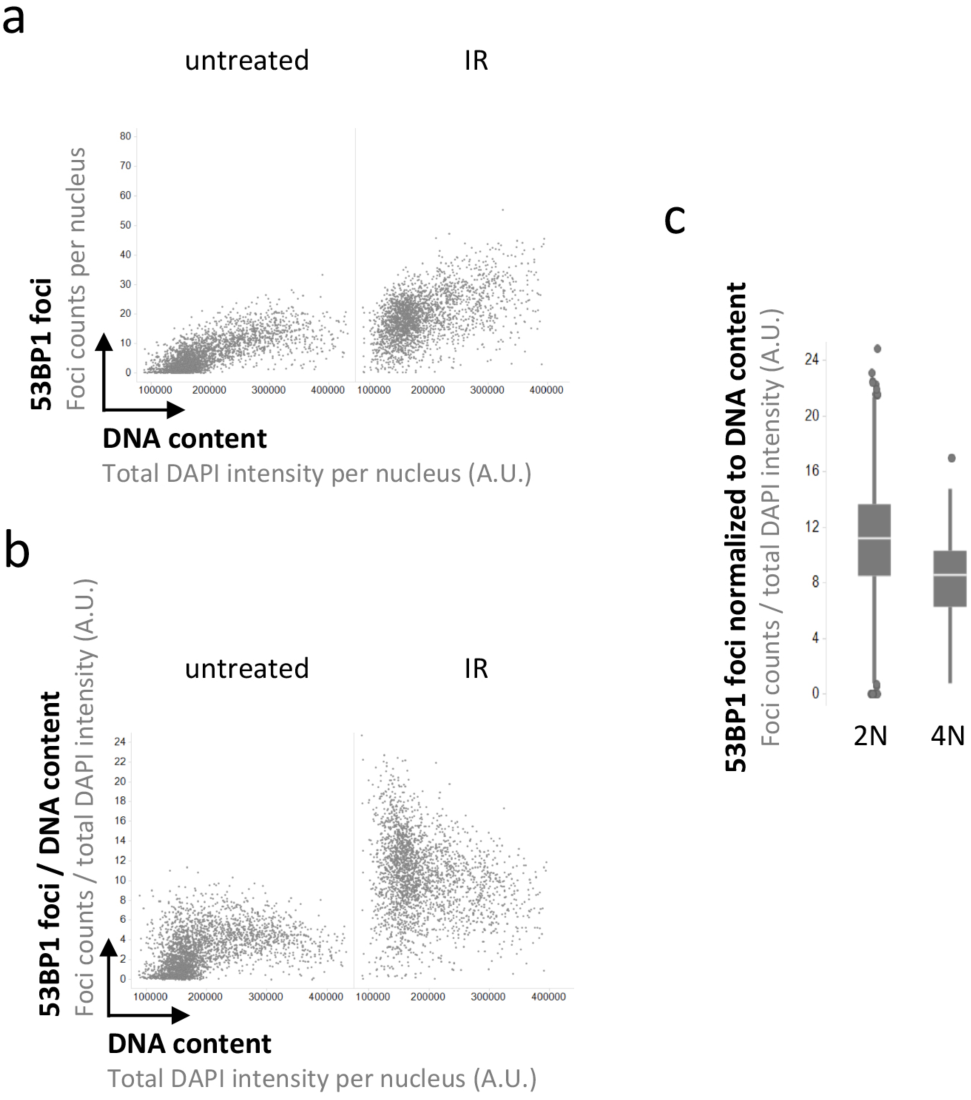
53BP1 recruitment to replicated chromatin is inefficient in *BRCA1*-mutated cancer cells. **(a)** Cell cycle-resolved analysis of IR-induced 53BP1 foci formation in *BRCA1*-mutant SUM149PT cells. SUM149PT cells were treated with 0.5 Gy of IR, fixed 45 minutes later, stained for DNA content and 53BP1, and 53BP1 foci were quantified by QIBC. **(b)** The quantification from (a) was normalized at the single cell level to DNA content to control for increasing damage load with increasing DNA amount. **(c)** 53BP1 recruitment from (b) is compared in 2N versus 4N cells. Box plots with medians are shown.

## Materials and Methods

### Cell culture and treatments

Human U-2 OS cells and U-2 OS cells expressing 53BP1-mScarlet from the endogenous promoter^25^ were grown under standard cell culture conditions (humidified atmosphere, 5% CO_2_) in Dulbecco’s modified Eagle’s medium (DMEM) containing 10% fetal bovine serum (GIBCO) and penicillin-streptomycin antibiotics. SUM149PT cells (kindly provided by Alessandro Sartori) were grown in Ham’s F-12 medium (Thermo Fisher Scientific) containing 10% fetal bovine serum and 1% of penicillin-streptomycin antibiotics. All cells were maintained in a sterile cell culture environment and routinely tested for mycoplasma contamination. Irradiation was performed with a Faxitron Cabinet X-ray System Model RX-650. Transfections with Ambion Silencer Select siRNAs were performed for 72h using Lipofectamine RNAiMAX (Thermo Fisher Scientific). The following Silencer Select siRNAs were used at a final siRNA concentration of 25nM: siBRCA1 (s459) and siBARD1 (s1887). Negative Silencer Select control Neg1 from Ambion was used as non-targeting control. For pulsed EdU (5-ethynyl-2’-desoxyuridine) (Thermo Fisher Scientific) incorporation, cells were incubated for 20 min in medium containing 10μM EdU. The Click-iT EdU Alexa Fluor Imaging Kit (Thermo Fisher Scientific) was used for EdU detection.

### Immunofluorescence

Cells were grown on sterile 12 mm glass coverslips, fixed in 3% formaldehyde in PBS for 15 min at room temperature, washed once in PBS, permeabilized for 5 min at room temperature in PBS supplemented with 0.2% Triton X-100 (Sigma-Aldrich), and washed twice in PBS. All primary and secondary antibodies (Alexa fluorophores, Life Technologies) were diluted in filtered DMEM containing 10% FBS and 0.02% Sodium Azide. Antibody incubations were performed for 2h at room temperature. Following antibody incubations, coverslips were washed once with PBS and incubated for 10 min with PBS containing 4’,6-Diamidino-2-Phenylindole Dihydrochloride (DAPI, 0.5μg/ml) at room temperature to stain DNA. After three washing steps in PBS, coverslips were briefly washed with distilled water and mounted on 6μl Mowiol-based mounting media. The following primary antibodies were used for immunostaining: H2AX Phospho S139 (mouse, Biolegend 613401, 1:1000), 53BP1 (mouse, Upstate MAB3802, 1:1000), H4K20me2 (rabbit, Abcam ab9052, 1:100), Cyclin A (mouse, Santa Cruz sc-271682, 1:100), and RAD51 (rabbit, Bioacademia 70-002, 1:1000).

### Quantitative image-based cytometry (QIBC)

Automated multichannel wide-field microscopy for quantitative image-based cytometry (QIBC) was performed on an Olympus ScanR Screening System equipped with an inverted motorized Olympus IX83 microscope, a motorized stage, IR-laser hardware autofocus, a fast emission filter wheel with single band emission filters, and a digital monochrome Hamamatsu ORCA-FLASH 4.0 V2 sCMOS camera (2048 × 2048 pixel, 12-bit dynamics). For each condition, image information of large cohorts of cells (typically at least 800 cells for the UPLSAPO 40x objective (NA 0.9), and at least 2000 cells for the UPLSAPO 20x objective (NA 0.75)) was acquired under non-saturating conditions at a single autofocus-directed z-position. Identical settings were applied to all samples within one experiment. Images were analyzed with the inbuilt Olympus ScanR Image Analysis Software Version 3.0.0, a dynamic background correction was applied, and detection of cell nuclei was performed using an integrated intensity-based object detection module based on the DAPI signal. All downstream analyses were focused on properly detected interphase nuclei containing a 2N-4N DNA content as measured by total and mean DAPI intensities. Fluorescence intensities were quantified and are depicted as arbitrary units. For normalization according to DNA content, measurement parameters (e.g. 53BP1 foci numbers) were divided at the single cell level by the DNA content (measured as total DAPI intensity per nucleus). The parameter resulting from this normalization has arbitrary units and was multiplied by a multiple of 10 to yield data that could be plotted on linear scale in the depicted range. Scatter plots of asynchronous cell populations were generated with Spotfire data visualization software (TIBCO). Within one experiment, similar cell numbers were compared for the different conditions. Representative scatter plots and quantifications of independent experiments, typically containing several thousand cells each, are shown.

## Acknowledgments

We are grateful to the University of Zurich Center for Microscopy and Image Analysis for microscopy support. We thank all members of our lab and of the DMMD and IMCR for discussions, M. Pruschy for input and advice on normalization of radiation damage, and A. Groth for helpful comments on the manuscript. Research in the lab of M.A. is supported by the Swiss National Science Foundation (grants 150690 and 179057), the European Research Council (ERC) under the European Union’s Horizon 2020 research and innovation program (ERC-2016-STG 714326) and the UZH Candoc/Postdoc program. The authors declare no competing interests.

## Author contributions

J.M., S.P., and V.S. designed and performed experiments and analyzed results. M.A. supervised the project and wrote the manuscript draft with input from all authors.

